# Disentangling eco-evolutionary effects on trait fixation

**DOI:** 10.1101/259069

**Authors:** Peter Czuppon, Chaitanya S. Gokhale

## Abstract

In population genetics, fixation of traits in a demographically changing population under frequency-independent selection has been extensively analysed. In evolutionary game theory, models of fixation have typically focused on fixed population sizes and frequency-dependent selection. A combination of demographic fluctuations with frequency-dependent interactions such as Lotka-Volterra dynamics has received comparatively little attention. We consider a stochastic, competitive Lotka-Volterra model with higher order interactions between two traits. The emerging individual based model allows for stochastic fluctuations in the frequencies of the two traits and the total population size. We calculate the fixation probability of a trait under differing competition coefficients. This fixation probability resembles qualitatively the deterministic evolutionary dynamics. Furthermore, we partially disentangle the selection effects into their ecological and evolutionary components. We find that changing the evolutionary selection strength also changes the population dynamics and vice versa. Thus, a clean separation of the ecological and evolutionary effects is not possible. The entangled eco-evolutionary processes thus cannot be ignored when determining fixation properties in a co-evolutionary system.

## 1 Introduction

The theoretical study of fixation or extinction of a trait in a population has a long history in the context of population genetics [Hal27, Fis22, Wri31, Kim62]. It has since then served as a basic theory in a huge variety of fields in evolutionary biology [NS13]. While traditional models of trait fixation consider fixed or infinitely large population sizes, also deterministically changing population sizes have received some attention [Ewe67, KO74, OW97, WG01, UH11].

Evolutionary game dynamics has been utilized in biological and social contexts since its inception [Lew61, MSP73]. This framework allows for an easy interpretation and implementation of frequency-dependent selection leading to coexistence or bistable dynamics. Evolutionary games have provided insights into evolution of cooperation [NSTF04], evolution of sex [MS86], host-parasite-dynamics [KS87] and more recently, evolution of cancer [PSD14]. The introduction of stochasticity in fixed population sizes, allows the study of quantities such as fixation probabilities or mean fixation times [GRD74]. The main focus of this discipline, has been on evolutionary dynamics [HS98, Now06], assuming ecological equilibrium.

In real biological systems, as evidenced from epidemiological and experimental studies, the interaction of ecology and evolution is crucial in determining the joint eco-evolutionary trajectory of a system [SG13, FSB16, HACB16]. Evolutionary dynamics of two traits, e.g. cooperators and cheaters, has been studied in evolutionary models inspired by microbial experiments [ASF+08, MFHG+10, CRL10]. Ecology, and in particular fluctuating population sizes, often dictate the dynamics of these experiments.

However, most theoretical studies focus on a constant population size, neglecting potential ecological effects on the population dynamics. Just recently the interaction of evolutionary and population dynamics has gained more attention [Lam06, CL07, PQ07a, PQ07b, PQP10, UH11, CM15, PR17]. Classical equations of ecology like the Lotka-Volterra dynamics have to be re-evaluated when finite populations are considered [GPTS13, PGTS16]. Additionally, recent studies explicitly include evolutionary game dynamics into an ecological framework [HHT15, GH16, ASGG17, YRD17, CT18, MFH+18]. In these models, one challenge is to re-interpret game interactions in terms of ecological dynamics to make sense in a fluctuating population size scenario. Since the game interactions are between individuals with different traits, we can interpret them as the interaction terms in the competitive Lotka-Volterra equations [Zee95, HHT15]. Typically in evolutionary games when an individual interacts with another, it receives a payoff. In our eco-evolutionary setting, the payoffs translate inversely into competition outcomes. Thus, the more the payoff, the less likely is the interaction harmful for the actor.

Derivation of a stochastic formulation of a model begs further analysis. When genetic drift dominates, i.e. in the limit of weak selection, the impact of interactions on fitness is minimal and approximations for the fixation probability are available [Lam06, CL07, CM15, CT18]. From a game theoretic perspective, all these studies are restricted to the highly abstract notion of two player games. The mathematics of these games is the same as that of allele dynamics within a haploid population [CK70, TH09]. This framework has been extended to diploids [Row88, HA09] and going further, multiplayer games would allow us to increase the ploidy level [HTG12]. Therefore, multiplayer games are not just theoretically interesting, but have clear biological as well as social interpretations. From multiple bacteria interacting together as in microbiomes [LPF+15, WR16], in quorum sensing [WSE17] or in biofilm formation [DNS+14] to social dilemmas such as the classic tragedy of the commons [Har68], multiplayer games can be interpreted across scales of organization. Of interest then, would be a complete eco-evolutionary analysis of fixation probabilities for multiplayer evolutionary games with demographic changes.

Following this line of thought, we develop an ecological interpretation of a two trait multiplayer evolutionary game. We calculate the fixation probability of a trait (strategy) in a competitive Lotka-Volterra model with higher order interactions (multiplayer game), where the population size fluctuates over time due to demographic noise. This individual based implementation of reactions allows for a straight-forward interpretation of fitness effects. The stochastic model so generated, generalises previous results on fixation probabilities in a similar setting [Lam06, CT18]. It allows us to (partly) disentangle the impact of evolutionary and ecological forces on the fixation probability. We then apply our theory to a well studied example of a social dilemma, the so called threshold public goods game. This example fleshes out the structure of the expression of the fixation probability allowing us to extend the framework to eco-evolutionary models with general *d*-player interactions.

## 2 Model and Methods

While two player games form the crux of most of evolutionary game theory, multiplayer games are rather the norm in social as well as a number of biological situations. Assuming population densities being in an ecological equilibrium the change in frequencies of traits can be calculated by the replicator equation. For changing population densities we develop a multiplayer population dynamics model which is based upon ecological processes as in [HHT15]. We begin with a three player interaction.

### 2.1 Replicator dynamics

Consider a population consisting of two traits, *A* and *B*. The interactions between individuals in groups of three are then denoted by,

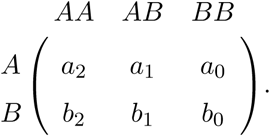

The focal individual (row) with trait *A* interacts with two other individuals. If the other two individuals happen to be *A* then the payoff to the focal individual is *a*_2_ and so forth. Typically assuming the population size to be infinitely large the number of *A* and *B* individuals can be represented by their frequencies *x*_*A*_ = *x* and *x*_*B*_ = 1 *– x*. The fitness of the trait is then the product of the payoff and the frequency of the corresponding trait in the population,

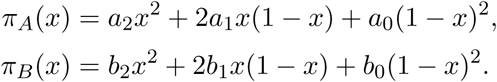

The evolutionary change in the frequencies of the traits can be captured by replicator dynamics, also valid for multiplayer games [HS98, GT10],

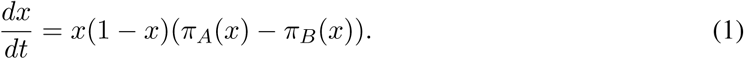

We can recover the traditional outcomes of neutrality, dominance, bistability and coexistence for the three player game. Furthermore the setup has the possibility to show two internal fixed points.

#### 2.1.1 Finite populations

Replicator dynamics allow us to look at the gross qualitative dynamics of selection. Interactions however take place in finite populations. Taking this into account forces us to rethink the limits of the infinite population size assumption [FAF98, TPI06]. There are various ways of handling finite populations, also in multiplayer games [TH09, GT10, Les11].

A crucial concept in finite populations is of selection intensity. We can control the effect of the game (interactions) on the fitness of a trait by tuning the magnitude of the intensity of selection. Assuming a linear payoff to fitness mapping we have *f*_*a*_ = 1 + *ωπ*_*A*_. If selection is weak, *ω ≪* 1, then genetic drift dominates and the average payoff has minimal effect on fitness. Thus the effective difference between the two traits reduces. The mapping can be subsumed in the payoff matrix where each payoff entry *a*_*i*_ is rescaled to 1 + *ωa*_*i*_.

### 2.2 Eco-evolutionary dynamics

As per [HHT15] we rationalize that since the game contributes only to competition between individuals, it can result in the death of the focal individual. This reflects the interpretation of individuals interacting with each other, similar to the game theoretic view on the payoff matrix. Additionally, this mapping from payoffs to population dynamics allows us to write the carrying capacities in terms of the payoff values, i.e. the single species equilibria are solely affected by *a*_2_ and *b*_0_, respectively, while the coexistence points are related to all the payoffs. Moreover, the payoffs of a game typically translate positively towards the fitness of the focal individual. Hence we assume an inverse relationship between the magnitude of the payoff and the death rate. The microscopic interactions which lead to birth and death of individuals can be written as follows

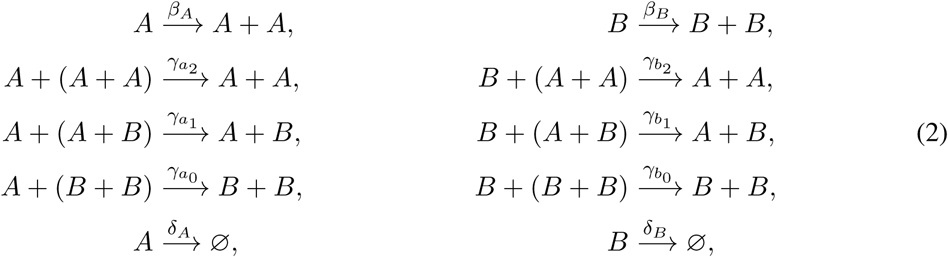

with reaction rates 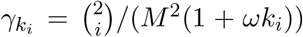. Here, *k*_*i*_ are the entries of the payoff matrix for the three player game and *M* is a scaling parameter describing the size of the population in equilibrium, i.e. the fixed points are proportional to *M*. The reactions are corrected according to their combinatorial possibilities, the factor 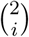, as done in [MN04]. The scaling *M*^2^ changes the absolute values of interacting individuals to their densities. This gives the right scaling for the population in stationarity to be proportional to *M*. It can also be interpreted in terms of the probability of meeting two individuals at the same time. Since our interactions happen amongst three individuals we need to scale these rates with *M* ^2^ rather than just *M* which would hold for binary interactions. For more details on the choice of scalings we refer to the literature concerned with chemical reactions, see for instance [Gil76, AK15].

Such a mechanistic implementation of evolutionary processes has not yet received much attention in the literature – see [DIS17] for an essay about this – but has the advantage of directly relating selective advantages to birth, death or competition processes. We believe that interpreting selection in this way can provide new insight into the concrete advantages of mutants and situations in which they are beneficial.

For realising our ultimate aim of approximating the fixation probability of trait *A* individuals in the population, we now consider the stochastic description of this eco-evolutionary model.

#### 2.2.1 Stochastic eco-evolutionary dynamics

Often stochastic models are derived from an individual based formulation as given in Eq. (2) and then approximated by a stochastic differential equation, see e.g. [vK97, Gar04]. Since we are mainly interested in the effect of individual interactions on the dynamics we set *β*_*A*_ = *β*_*B*_ = *β* and *δ*_*A*_ = *δ*_*B*_ = *δ*. Further, denoting by *X*_*i*_ the absolute number of type *i* individuals and setting *x*_*i*_ = *X*_*i*_*/M* the corresponding density we obtain the following stochastic equations of our system (see A for a detailed derivation)

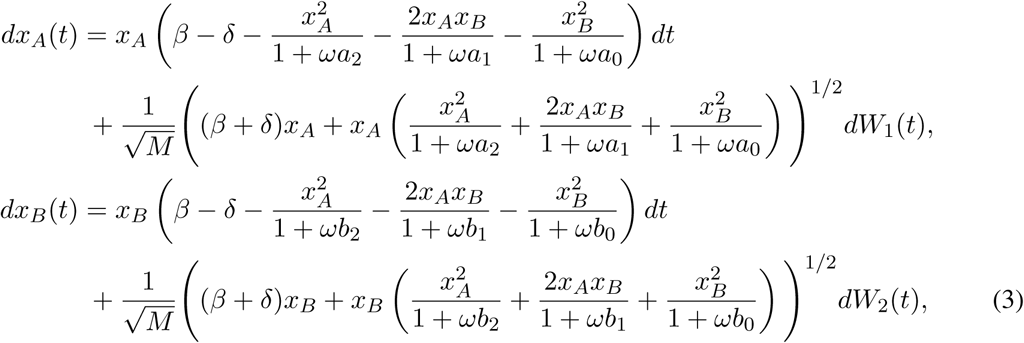

where *W*_*i*_ are independent Brownian motions. We can describe the evolutionary process (change in the population composition) as well as the ecological dynamics (change in population density) by transforming the system to the variables, 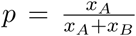 the frequency of trait *A* individuals and *z* = *x*_*A*_ + *x*_*B*_, the total population density, see also Figure 1. The figure illustrates the transformation from the (*x*_*A*_, *x*_*B*_)-space (bottom left) to the (*p, z*)-space (bottom right). Additionally, we show the replicator dynamics at the right. There, the deterministic trajectories are attracted to the stable equilibrium (filled circle) while escaping the unstable equilibrium (open circle). In general a *d–*player interaction allows for *d–* 1 internal equilibria in a Lotka-Volterra system. This can be derived by solving the deterministic part of equation (3). Furthermore, the stability of the internal equilibria is alternating. In our example in Figure 1, 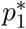 is stable and 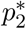 is unstable. Another interesting feature in our system is that the dynamics quickly collapse on the ecological equilibrium due to the Lotka-Volterra dynamics. Systems where the ecological component of the system is slow and the evolutionary dynamics evolve on a faster time-scale are for instance analysed in [PCS18].

**Figure 1:**
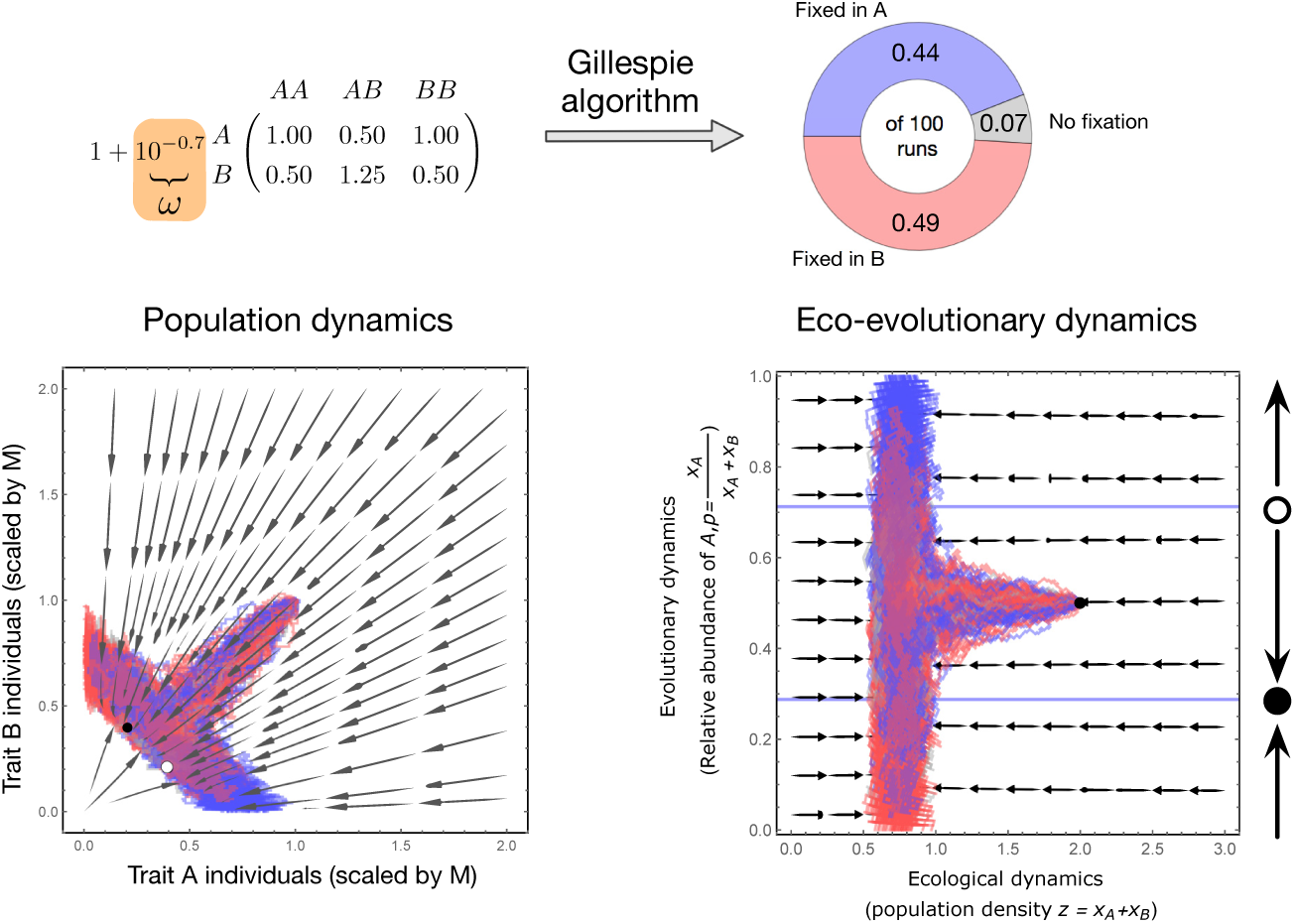
From an evolutionary game to population dynamics. The interaction between traits *A* and *B* is represented by the payoff matrix as shown in the top left of the figure. We include weak selection via a linear payoff to fitness mapping such that the effective matrix is 1 + *ω ×* matrix. Then we compute the population dynamics of this interaction matrix for weak selection *ω* = 10^*–*0.7^. Thus, as the dynamics comes close to neutrality the number of runs fixed in either trait *A* or *B* is almost equal. From an eco-evolutionary point of view (bottom right), the population density rapidly converges to the ecological equilibrium and then the almost neutral dynamics proceeds in the evolutionary dimension of the fraction of trait *A* individuals. The fixed points in the evolutionary dimension are denoted by the solid horizontal lines (given explicitly in the Appendix) and under weak selection coincide with the equilibria of the deterministic replicator equation. The replicator dynamics are illustrated at the right of this subfigure, showing that trajectories point away from the unstable fixed point (open circle) towards fixation or the stable fixed point (filled bullet).

## 3 Results

### 3.1 Fixation Probability

Under weak selection we are able to approximate the fixation probability *ϕ*(*p*_0_, *z*_0_) of trait *A*. It depends only on the initial population size *z*_0_ and the composition of the initial population defined by *p*_0_. The techniques we use to derive an interpretable expression are first described in [Lam06] and [CL07] and refined for this specific setup in [CT18]. The interpretation relies on our ability to separate the evolutionary terms from the ecological variable *z*. For two player games, the condition for this sep-aration to be valid coincides with the weak selection assumption [CT18]. For multiplayer games, the separation of evolutionary and ecological scales becomes more complicated and harder to interpret. By choosing a linear payoff function 1 + (*ω × ·*), we are able to split weak selection into its evolutionary and ecological components. The approximation further depends on the number of internal equilibria. A three player interaction can have 0 *–* 2 internal fixed points dependent on the payoff matrix associated with these interactions. We first consider the situation with two internal fixed points since it is the most insightful for understanding the potential and limitation of the approximation. Denoting by 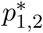 the equilibria of the deterministic model in the *p*-space, we find the following conditions under which we can separate the evolutionary from the ecological scale:

(i) ω≪1,

(ii) 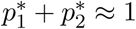

(iii) 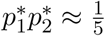

Condition (i) is the overall impact of the trait differences on the population’s development – the evolutionary component. Condition (ii) and (iii) ensure that selection is negligible over the whole frequency space, i.e. the integral of the replicator dynamics over the frequency space is zero (or close to it). Formally, conditions (ii) and (iii) yield

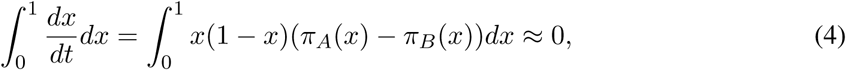

where the equality is explained by Eq. (1). This is an extension of the two player case where the ecological condition for the derivation of the corresponding result is *p*^*\**^ *≈* 1*/*2 [CT18]. Thus conditions (ii) and (iii) form – the ecological component – of weak selection. Furthermore, they impose the payoff matrix to allow for two internal equilibria since solving for 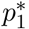 and 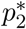 here yields: 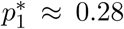 and 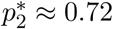.

Conditions (ii) and (iii) seem to be very restrictive, however they emerge naturally if we impose that mutations are close to neutral, i.e. trait *A* and *B* are similar. Weak mutations (or nearly neutral mutations) can be translated to having negligible selective (dis-)advantage which can be related to equation (4) being close to zero.

Another approach to analyse the system above is by explicitly separating the time-scales such that the system reduces to a one-dimensional stochastic diffusion process on a manifold where population size is fixed. H owever, t his t ime-scale s eparation i s o nly v alid u nder w eak s election s uch t hat we suspect similar conditions to be found to (i)-(iii) for the technique to work. A nice review of methods using this approach is [PR17].

Taken together, conditions (i)-(iii) already show that weak selection in an eco-evolutionary model can arise from multiple sources. Either, (a) the whole model follows weak selection, i.e. all conditions are satisfied a t t he s ame t ime o r (b) a m odel c an b e s tudied u nder e cological o r evolutionary weak selection. We analyse the different impacts of the above assumptions on both such situations.

Satisfying all conditions, we can derive the following expression for the fixation probability (details of the derivation are stated in B):

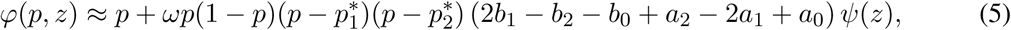

where *ψ*(*z*) is a function indepe*\**1ndent of *p* satisfying a differential equation explicitly given in B. Note that all the ecological information is gathered in the function *ψ*(*z*). This separates the evolutionary and ecological impact on the fixation probability under weak selection and nearly neutral mutations. Comparing this result to similar results from the population genetics literature the function *ψ*(*z*), as a measure of impact of the initial population size, takes the place of the effective population size in related formulas. This seems reasonable since it is independent of the initial trait frequencies and therefore only dependent on the population dynamics.

The approximation shows a good fit with results obtained from stochastic simulations with parameters satisfying conditions (i)-(iii), see Fig. 2 (dotted line). For a sensitivity analysis we refer to Section 3.3.

**Figure 2:**
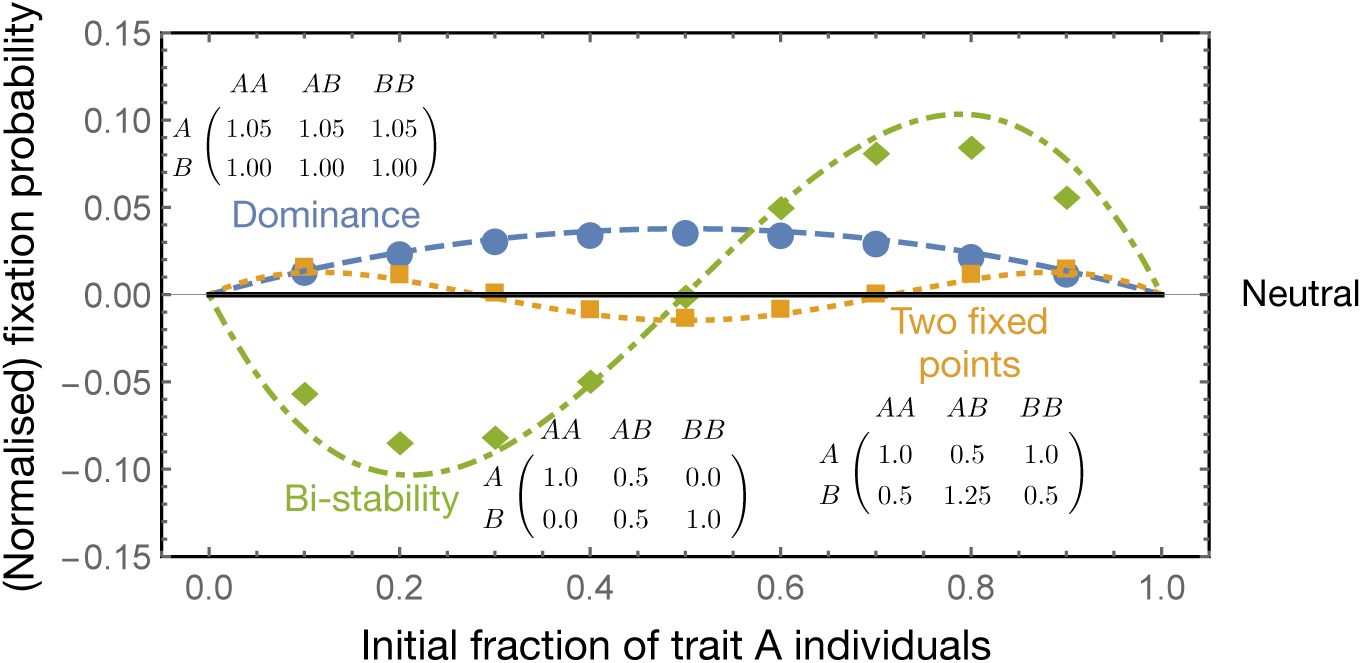
Generality of our approach encompassing all possible three player outcomes. Linear (two player) evolutionary games can result in neutral, dominance (no internal fixed points) or co-existence and bistable (one-interior fixed point) dynamics. In addition to these, a three player game allows for two interior fixed points. We show that our analysis i s able to capture the fixation probability not just in the scenarios reminiscent of linear (two player) games but encompass the whole spectrum of outcomes possible in non-linear (three player) games. The fixation probability (*y*-axis) is normalised in each case by the neutral expectation (by subtracting *p*, the initial fraction of trait *A* individuals *x*-axis). The payoff matrices for the dominance, bistability and two fixed points scenarios are as stated in the p lot. Symbols are simulation results, averaged over 10^6^ realisations. The theoretical results for the three cases are given by Eqs. (5),(6) and (7) for two internal fixed points (dotted), bistability (dot-dashed) and dominance (dashed), respectively (with *ψ*(*z*) being 5.8486, 10.7398 and 30.1971, respectively). The other parameters are fixed in all scenarios and set to *M* = 100, *β* = 0.6, δ = 0.1 and initial population size of 200.

### 3.2 Generality of the approximation technique

The previous derivation holds in the case of two internal fixed points and for interaction rates that allow for these equilibria to satisfy conditions (ii) and (iii).

Assuming exactly one internal fixed p oint t he a nalysis b ecomes m ore i nvolved s ince t he scale separation does not follow as intuitively as in the previous case. Indeed, we find that an approximation is only possible in case the payoff values satisfy

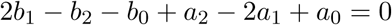

and in addition allow for an internal fixed point. The crucial part is the separation of the variables *p* and *z* that is ensured by the stated condition. It turns out that parameter configurations like this produce an internal fixed point close to 1*/*2, reminiscent of the analysis in [CT18], in fact the fixation probability takes the exact same form:

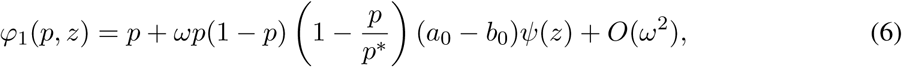

where the fixed point is given by

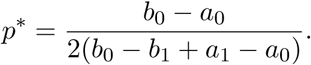

However, the partial differential equation*ψ* (*z*) needs to solve differs slightly from the two player scenario. The equation is stated, along with the derivation of the fixation probability, in B.1.

At last we consider the scenario with a purely dominant strategy, i.e. *a*_*i*_ *> b*_*i*_ or *b*_*i*_ *> a*_*i*_ for all *i ∈* {0, 1, 2}. In this case, in addition to condition (i) we need the payoff values to be close to each other, similar to the fact that the selective advantage of one type needs to be negligible over the frequency space, see also equation (4). To be more precise we need the following products to be vanishingly small:

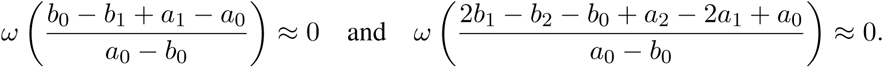

Then the fixation probability can be approximated by

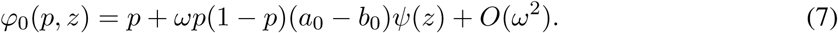

The derivation is the same as in the corresponding two player scenario [CT18, Theorem 2], again with a different partial differential equation for *ψ*(*z*), but is stated for completeness in B.2.

Choosing parameter values that satisfy the conditions necessary for these approximations to work, we see that indeed the calculated fixation probability fits the simulated data, see Fig. 2 the solid and the dash-dotted line for the dominance and bistable case, respectively.

### 3.3 Breaking assumptions in the two internal fixed points scenario

In the following we examine the impact of deviating from conditions (i)-(iii) under which the theory developed for two internal fixed points holds. Therefore, as a “benchmark” model we choose the matrix

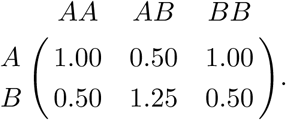

The internal equilibria corresponding to this matrix are given by 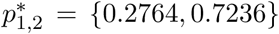. For this combination we have 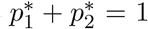 and 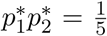 which holds for *ω ≪* 1 thus satisfying the required conditions.

As already mentioned our setting allows for two possible ways to deviate from the idea of weak selection. We can assume larger values of *ω* corresponding to an overall larger impact of the evolutionary fitness differences on the model. By increasing *ω* the fixation probability is less and less captured by our theoretical expectation, see *x*-axis of Fig. 3. Alternatively, weak selection is achieved when the payoff values of the matrix satisfy conditions (ii) and (iii) from above. Varying the payoffs changes the intensity of selection but more importantly also alters the location of the fixed points. In this case the theory is able to capture qualitatively the direction of deviation from neutrality (Fig. 3 *y*-axis). We call these two deviations either evolutionary (*x*-axis) or ecological (*y*-axis) weak selection even though it is not possible to strictly disentangle the effects as they are intertwined via the calculation of the fixed points.

**Figure 3:**
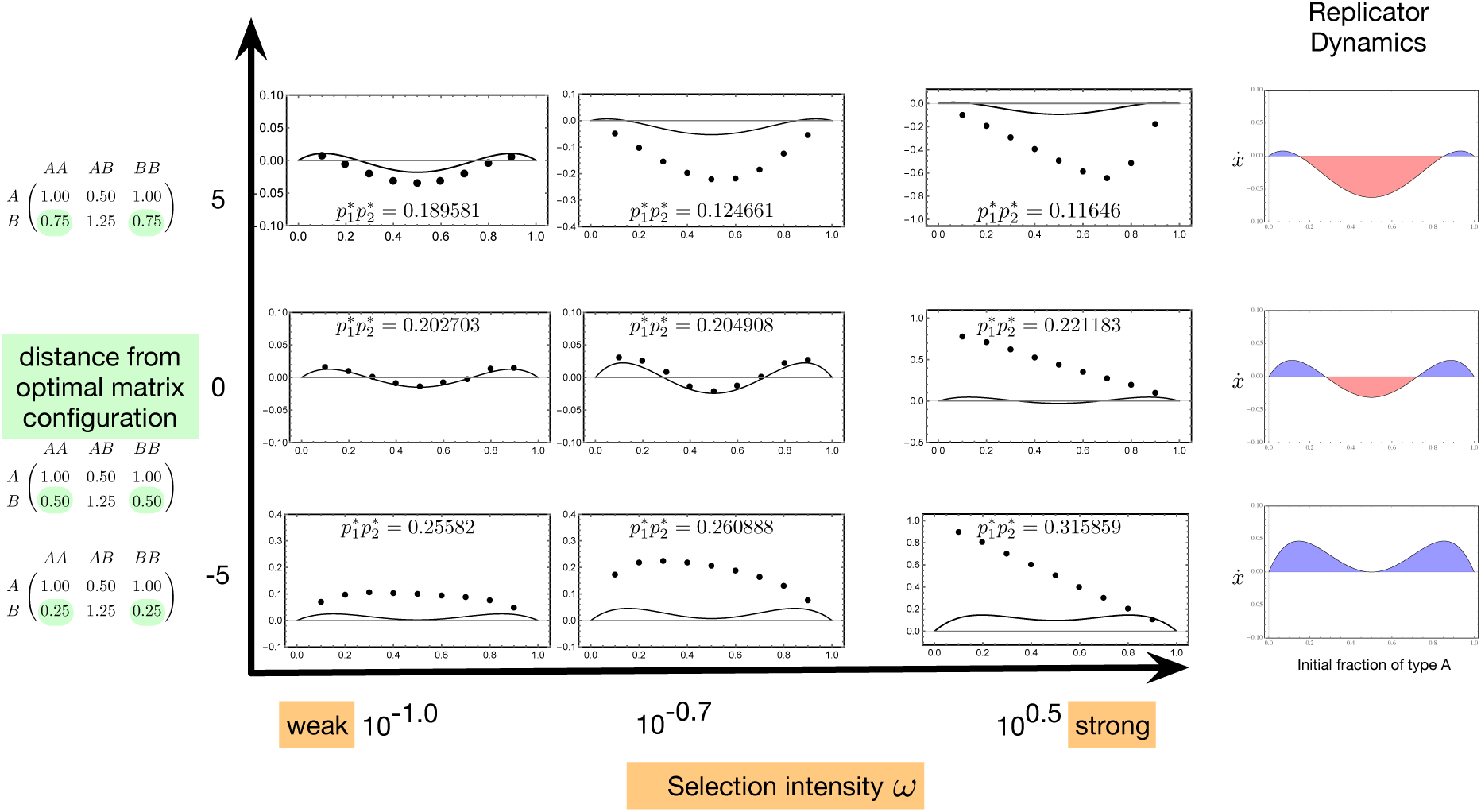
Weak selection(s). Typically in evolutionary games, scaling the payoffs (by the selection intensity) does not change the deterministic dynamics. Thus across the *x*-axis the replicator dynamics Eq. (1) can be visualized, see right column. A positive value (blue) determines that trait *A* is favoured over trait *B* and vice versa for negative values (red). In our setting the intensity of selection plays a major role (*x*-axis). For weaker selection, the magnitudes of the positive and negative values are extremely small, tending towards neutrality. Taking population dynamics into account, another way of introducing weak selection is when the payoff entries are close to our “benchmark” model (*y*-axis). If we increase the difference between the payoffs, we change the dynamics of the game and the fixed points of the (ecological) system, thus also affecting the regions where the theory is applicable 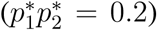. One unit of change on the *y*-axis corresponds to a change of 0.05 in the highlighted payoffs. As the payoffs change the fixed points move symmetrically towards each other (for a negative change) and away from each other (for a positive change). The insets show fixation probabilities of the stated payoff matrices and shown *ω* values as in Fig. 2. The simulations, depicted as dots, were averaged over 10^5^ realizations. For the cases where the fixed points become imaginary (bottom line, mid and right inset), the product of the fixed points is the sum of the products of the real and imaginary parts.

#### 3.3.1 Evolutionary weak selection - varying *ω*

Typically in evolutionary games, changing the intensity of selection does not affect the fixed points in the infinite population size limit. This only holds approximately in our eco-evolutionary model. Due to varying the influence of the underlying game on the competition parameters we change the location of the fixed points in the system. Calculating fitness as 1 + *ω*(*matrix*), weak selection can be imposed when *ω ≪* 1. In this case we get the theoretical optimum value for 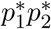 (Fig. 3, 4 horizontal axis). Increasing the values of *ω* leads to a deviation of 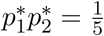 and thus the approximations become worse (Fig. 3). However, Figure 4 shows that we can recover this formally strict separation of *p* and *z* by moving along the *y*-axis, i.e. changes in the payoff matrix, such that even for higher values of *ω* the predicted fixation probability fits well to the observed values obtained by simulations (see the set of results for value 1 on the *y*-axis in Fig. 4).

**Figure 4:**
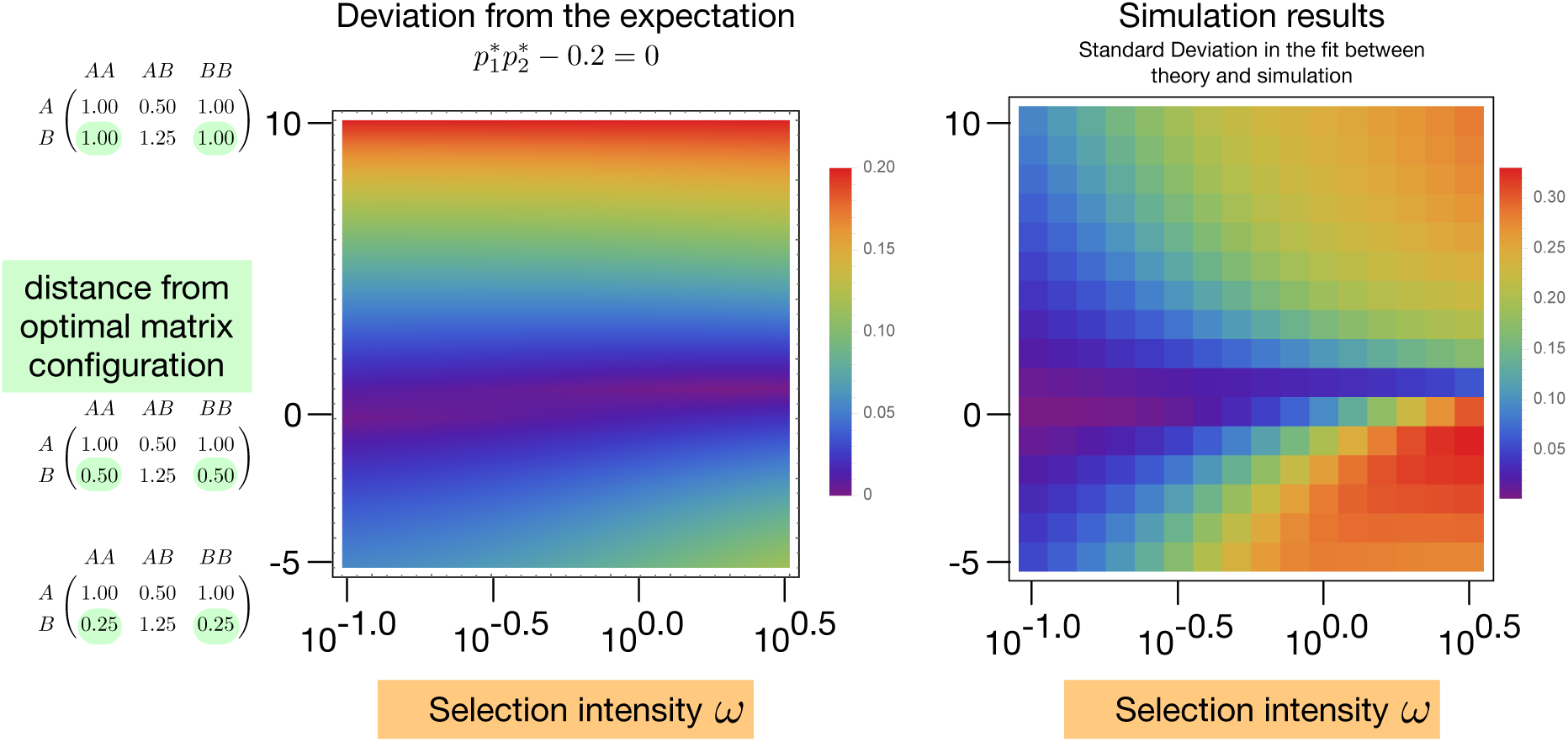
Studying weak selection. As we change the payoff entries in steps of 0.05 *× y* we are changing the fixed points such that 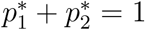 still holds but the condition of 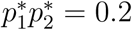 is not necessarily met (*y*-axis). In contrast we can modify *ω* which keeps the fixed points of the replicator dynamics constant but alters the fixed points of the corresponding Lotka-Volterra model. It does so by varying the interaction strength between the strategies (*x*-axis). As we change these two selection intensities, the expected theoretical performance under a strict separation of the variables *p* and *z* is illustrated in the left panel as the magnitude by which the condition 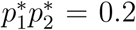 is violated. Gillespie simulations, starting at different initial fractions of *A* individuals (0.1, 0.2,…, 0.9) were performed and the fixation probabilities were calculated over 10^5^ realizations. The total initial population size was set to 200 with *M* = 100, *β* = 0.6, *δ* = 0.1 resulting in *ψ ≈* 5.85. We calculate the mean standard deviation between the simulation results and the expectation from Eq. (5). The right panel is a heat map of such deviations from the expectation for different *ω* and different matrix configurations. We see that the deviation is the least where the violation of 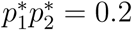 is the least.

#### 3.3.2 Ecological weak selection - varying the payoffs

When altering the payoffs, we change the location of the equilibria drastically which can be seen as changing the ecological output of the model, i.e. the carrying capacities of the two strains. For instance, in the monomorphic states these are given by 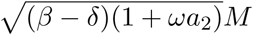 and 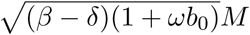 for trait *A* and *B*, respectively. Fixing *ω* and changing the payoff values indeed only affects the ecological dynamics since the impact itself on the evolutionary level is determined by *ω* and thus constant. Hence, to implement weak selection on the ecological scale, i.e. breaking assumptions (ii) and (iii) from above, we change the payoff entries keeping 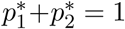 but violating 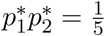 (Fig. 4 vertical axis). In a three player game a necessary condition for observing two internal fixed points is having two sign changes in *a*_*i*_ *– b*_*i*_ [HMND06, GT10, HTG12]. Since the matrix that we use is symmetric in the magnitude of the difference, the fixed points are also symmetric and hence the first assumption is maintained whenever the fixed points exist. The magnitude of the difference affects how close we get to satisfying the second assumption. Thus we modify only two payoff entries in the matrix, as shown on the *y*-axis in Fig. 3. The payoff matrix at *y* = 0 satisfies the condition 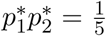 perfectly. As we increase the value of the highlighted payoff entries, the fixed points move closer to the edge in frequency space *p* with 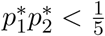.

For a decreasing value of the entries we have the fixed points colliding and eventually annihilating each other with 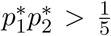. In case of vanishing fixed points, they become imaginary and their product is given by the sum of the products of real and imaginary parts.

Interestingly, for large selection intensities *ω*, the goodness of fit is non-monotonic for stronger deviations of condition (iii) (last column in Fig. 4). The discrepancy that we see for high *ω* and negative *y*-axis values can be attributed to ecological shifts of the equilibria. In the region of the discrepancy, the theory estimates the fixation probability still in the shape of the replicator dynamics, which is a gross underestimate (for example *ω* = 10^0.5^ and distance = 0 in Fig. 3.) As the distance decreases further, the replicator dynamics and the fixation probability both, increase above neutrality, reducing the standard deviation from the simulation results (*ω* = 10^0.5^ and distance = *–*5). No such discrepancy exists for positive *y* values since the simulations are progressively overestimated while maintaining the qualitative picture (*ω* = 10^0.5^ and distance = 5).

Overall, our analysis shows that the approximation provides a good fit in case all the assumptions are satisfied. Varying the payoff values has a large impact on the location of the ecological equilibria causing the prediction to deviate from the simulated data. On the other hand changing values for *ω* controls the effect the competition rates have, i.e. for *ω* = 0 the payoffs do not affect the model dynamics while for larger *ω*, the interaction rates have a large impact on the system. Hence, one should think of a nested selection characterization (evolutionary impact regulating the ecological variance) rather than two distinct variables acting on different scales. This entanglement is also highlighted in Figure 4 where the left heat map shows the expected outcome under a clean scale separation while the right heat map, comparing the actual simulation results to the prediction illustrates a complex interaction between the ecological and evolutionary scale.

### 3.4 Population dynamics of collective action

We now extend the analytical calculations to a particular example of multiplayer game. The evolutionary dynamics of collective action is an extremely well studied topic in the social sciences [Ost90, OBF+99]. How social structures overcome the tragedy of the commons is a recurring theme in this field [Har68, Sky03]. The tragedy of the commons is a case where the defectors benefit at a cost to the cooperators. However the tragedy is relaxed if a part of the benefit can be recovered by the acting cooperator. This negative frequency dependence is the essence of the snowdrift game [DHK04]. It has been proposed that the snowdrift game might better reflect human social dilemmas than the otherwise famous Prisoner’s Dilemma [KCF+07]. Also biological observations like phenotypic heterogeneity, a well established phenomena in microbes, can be a result of snowdrift like, negative frequency dependent dynamics [HAG16].

#### 3.4.1 (Multiplayer) Snowdrift game

The metaphor (for *d* players) states that *d* drivers meet at an intersection where a snowdrift has occurred and need to clear the snow in order to go home. If *k* drivers shovel they receive *b – c/k* (*c* being the cost of cooperating) while defectors get home for free, obtaining the pure benefit *b*. We are interested in further realistic cases where a certain threshold number of cooperators is needed to clear the street [SPS09]. Hence if *θ* is the threshold number of cooperators *necessary* to generate the benefit, then for *k < θ* the efforts of the cooperators go waste (0 *– c/θ*) while defectors receive nothing. This assumes that when the number of cooperators is less than the threshold value, cooperators still try to “give their best” as in [SPS09]. Once the number of cooperators is more than or equal to *θ*, the road is cleared. Cooperators, having paid the cost, get *b – c/k* while defectors receive *b*. If the number of other cooperators is *θ –* 1 then it is profitable to be a cooperator, fill the quorum, and reap the benefit *b – c/θ* as opposed to defect and end up with nothing. This concept of threshold public goods games is applicable not just in humans but in other species as well [Sta92, AN02, Lyo07, AMFC12].

Excluding finite populations precludes the possibility of observing ecologically relevant events such as extinctions. Population dynamics in social dilemmas have been considered before via deterministic dynamics [HHD06, GH16]. In a stochastic system as ours, the fixed points 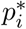 for a multiplayer game in the eco-evolutionary space (*p, z*) can be calculated in a similar way as in the three player game from above, see also B.3. In general for any *d–*player game the fixation probability can then be approximated by (details are stated in B.3)

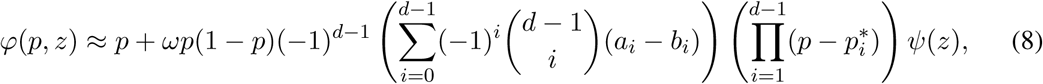

where *ψ*(*z*) is again given as the solution of a certain partial differential equation. We note that Eq. (8) reduces to the already obtained fixation probabilities in the cases *d* = 2, see [CT18, Theorem 1], and *d* = 3, see Eq. (5). The main features of the fixation probability are the same as in the cases with less players (intersections with the neutral fixation probability at internal equilibria, qualitative agreement with the replicator dynamics, i.e. *ϕ*(*p, z*) *– p >* 0 whenever 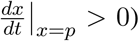. Exceptionally, the general approximation makes clear that the internal (and meaningful) fixed points determine the evolutionary success together with the other solutions of the corresponding deterministic model.

Despite uncertainties about the exact conditions for the approximation in (8) to be valid, it still captures the qualitative behaviour of the fixation probability of the system, reflecting the replicator dynamics (Fig. 5).

**Figure 5:**
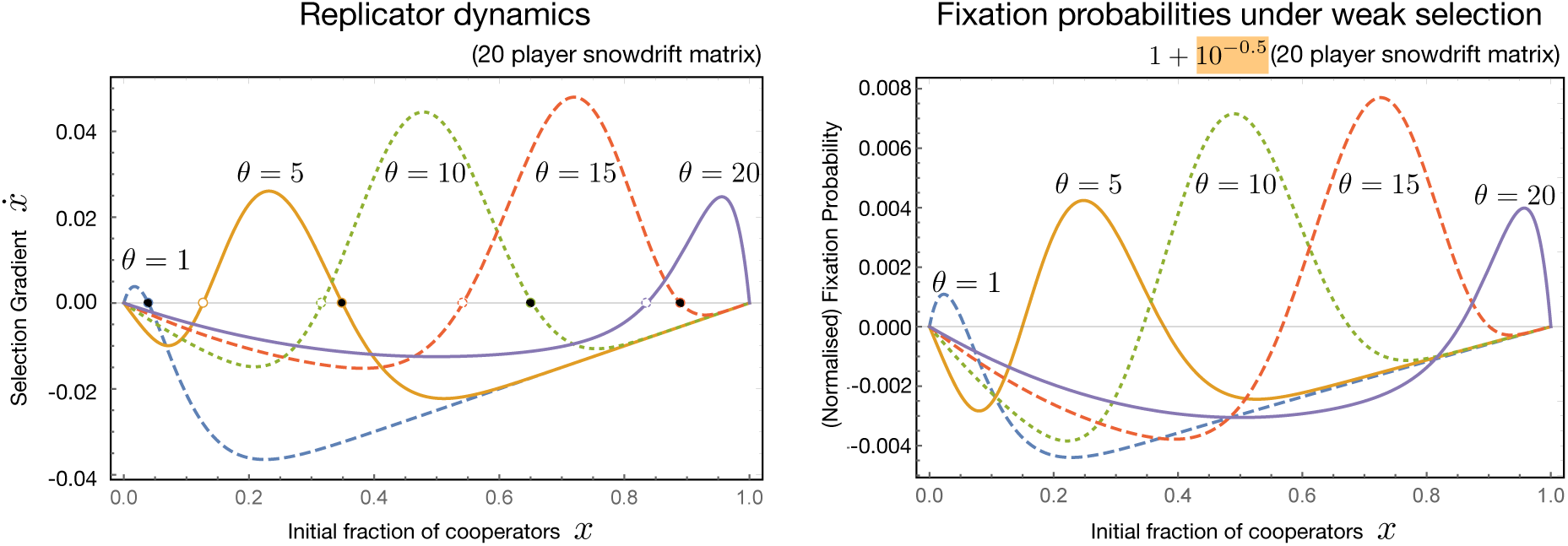
Fixation probabilities in a snowdrift game. The graph (on the left) shows the evolutionary dynamics as per the replicator equation for a 20 player game with different number of threshold snowdrift game scenarios (*θ* = 1, 5, 10, 15 and 20), benefit *b* = 1.5 and cost *c* = 1. For each case we calculate the corresponding fixation probability using Eq. (8) (right panel) for *M* = 100, *β* = 0.6, *δ* = 0.1, initial total population size of 200, weak selection *ω* = 10^*–*0.5^ and *ψ ≈* 0.53 (even though the parameters do not satisfy the assumptions on the separation of the scales). Comparing the structure of the fixation probability we see that it follows the gradient of selection qualitatively.

Varying the threshold number of necessary cooperators (*θ*) in a snowdrift game with *d* = 20 players we change the location of the two internal fixed points in the replicator dynamics [SPS09] (see left panel in Fig. 5). Setting *θ* = 10 and starting with equal numbers of cooperators and defectors, we estimate the fixation probability from stochastic simulations while varying *ω* (Fig. 6). For weak selection the trajectories almost randomly fix in either *allC* or *allD*. As selection increases, the distance between the two fixed points starts to matter. For example, for *ω* = 10^0.1^ the fixed points within the relevant space are 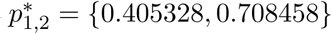. It is closer for the trajectories to go from 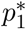 to the stable fixed point 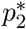 (and thus to *allC*) than to *allD*, hence increasing the likelihood of reaching *allC* (see intermediate *ω* in Fig. 6). Interestingly, for the results with strong selection stochastic drift may be an explanation. We find that the hitting probability of a Brownian motion gives a very accurate approximation of the fixation probability in case of strong selection, see B.5 for details.

**Figure 6:**
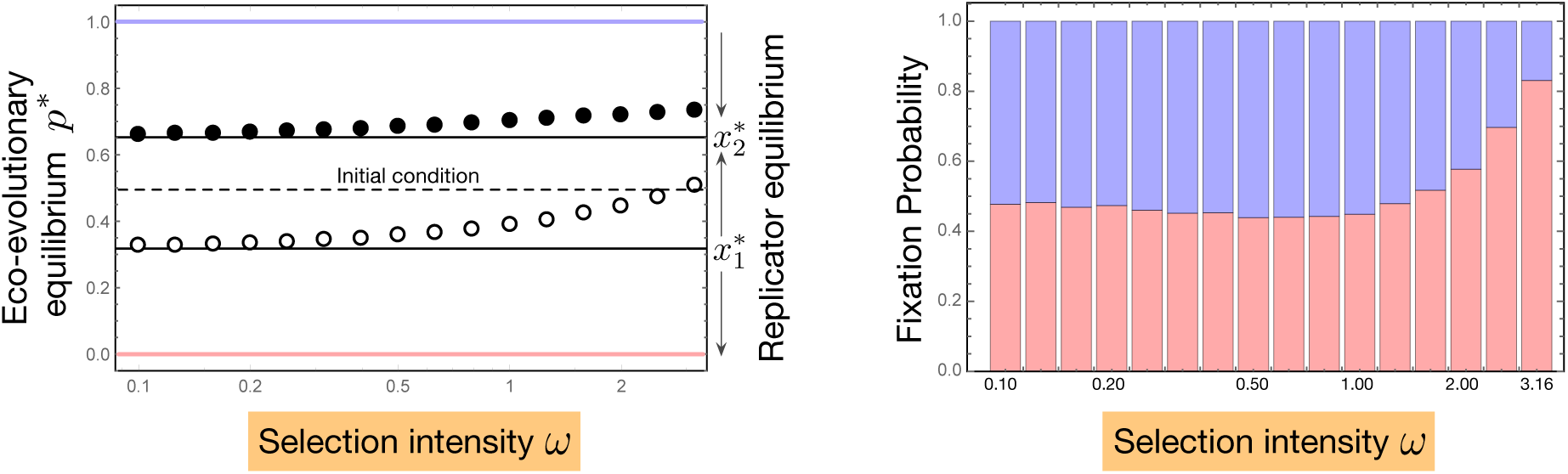
Snowdrift under selection. The threshold snowdrift game can have two internal equilibria 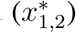 depending on the value of *θ*. In a finite population fixation occurs in either *allC* (blue, *p*^*\**^ = 1) or *allD* (red, *p*^*\**^ = 0). For weak selection the effect of the selection gradient is minimal and genetic drift plays a dominant role resulting in fixation probabilities close to neutral. As selection increases, the stable fixed point (filled circle), close to *allC*, attracts more trajectories which increases the probability of fixing in that state. However, for extremely strong selection the stable point is strongly attracting and fixation is again explained by random drift. Either the population fixes in the *allC* state or it overcomes the unstable point (empty circle) and then selection drives the population to *allD*. We performed 10^5^ Gillespie simulations for *M* = 100 and a 20 player snowdrift game with an initial population of 200, *b* = 1.5, *c* = 1, *β* = 0.6, *δ* = 0.1, and 1 + *ω*(*matrix*) mapping for different intensities of selection with *ω* = 10^*–*1.0^,…, 10^0.5^. The bar charts show the probability of a trajectory fixing in *allC* (blue) or *allD* (red).

## 4 Discussion

In co-evolutionary systems, populations are repeatedly subjected to changes in population sizes. Typical host-parasite systems undergo periodic cycles, where the population sizes are actively regulated by the antagonist. Early on, it was shown how the population size of the azuki bean weevil was affected by its parasitoid wasp [Uti57]. However the drastic population size change can affect other important evolutionary properties such as genetic diversity and evolutionary selection on traits [OW97, PQP10, HSD+14].

Understanding the fixation behaviour of traits in complicated situations where multiple individuals interact with each other is therefore relevant for understanding complex fixation patterns as for instance observed in [SG13, ML15]. Providing analytical insight into these processes offers theoretical explanations for non-trivial fixation behaviour. Extending [CT18] we have derived the fixation probability of a new trait in a stochastic Lotka-Volterra-model including higher order dynamics. These higher order interactions can be interpreted in terms of multiplayer games from evolutionary game theory or, as recently done in [MCA18] as mutualistic interactions. We explicitly deal with finite, fluctuating population sizes which are, for example, important in the context of co-evolution [Zee95, GPTS13, PK09, SGP+15]. While constant or infinite population size is captured by birth-death processes and the replicator equation, we apply stochastic diffusion theory to tackle demographic fluctuations.

Going from two player to three player interactions might seem like a minor extension but often, two body interactions are a special case. The extension to multiple players indeed helps us to see a general outline of the assumptions underlying the theory. While two player games required the internal fixed point to be at 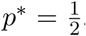, the conditions for higher order games clarify that in fact the requirement is that the fitness of the strategies over the complete frequency space needs to be balanced. More precisely, conditions (ii) and (iii), i.e. 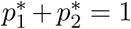 and 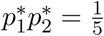, ensure that the selective advantage in the evolutionary dynamics of the replicator equation vanishes when considered over the whole frequency space. This linkage between the deterministic equilibria of Lotka-Volterra dynamics and the trait frequency dynamics highlight the fact that the intertwined nature of ecological and evolutionary effects cannot be entirely separated (see also Figures 3 and 4). Still, we are able to compute an explicit expression for the fixation probability at the cost of being restricted to certain parameter sets. Applying different techniques of scale separation, like for instance outlined in [PR17], could possibly help in extending the accessible parameter range. However, this type of analysis is beyond the scope of our study.

Besides providing a generalisation of the fixation probability for multiplayer games in populations of fluctuating size, we apply our analysis in the context of social evolution. Particularly we focus on the threshold version of social dilemmas, resonating with the concept of quorum sensing in microbes. Assuming that microbes play a two player game, i.e. linear interactions, is an assumption which can be easily violated in general biological systems [ST78, LPF+15]. In terms of Lotka-Volterra dynamics, this translates into including non-linear interaction terms. For typical predator prey dynamics, interactions can be captured by Holling response which provides a functional form to the interactions [Hol59]. However if the prey exhibits group defence or other forms of density depending modulation of the response we need to take the non-linearity into account [FHM86]. We focus on the first step towards this approach by studying a three player game. This small extension already results in mathematical complications which are non-trivial to analyse. For a general multiplayer game, we show that it is not easily possible to extrapolate the quantitative analysis of the fixation probability from the three player case. The reason, again, is that the separation of scales for high dimensional interactions is complicated to justify. We suspect that for a *d* player game, *d –* 1 different conditions would need to be satisfied.

Still, the replicator dynamics already allows us to estimate the qualitative structure of the fixation probability (Fig. 5) when including Lotka-Volterra type interactions. This alignment of deterministic and stochastic predictions is not a general rule as studies with fixed population sizes calculating fixation probabilities show distinctly opposite qualitative behaviour. A well known example of this phenomena is the one-third rule where a deterministically unfavourable trait can have a larger than neutral fixation probability [NSTF04, OBN07].

## 5 Conclusion

In conclusion, we have brought together the concept of weak selection from population genetics, multiplayer games from evolutionary game theory and population dynamics from theoretical ecology. By the synthesis of these fields, we have added new insight on the dynamics of fixation under demographic fluctuations by extending the competitive Lotka-Volterra model to now include higher order interactions between traits. The generality of our results comes from including higher order interactions but they are restrictive when it comes to the evolutionary impact of trait differences. Conditions (i)-(iii) only allow studying systems under weak selection in both, the evolutionary and the ecological, components. We observe that by increasing the complexity of the model, the separation of ecological and evolutionary processes becomes more and more difficult. For our specific purpose we find the influence of eco-evolutionary feedbacks to be crucial. In general, this eco-evolutionary feedback can have fateful impacts on the existence of communities of interacting species [PCS18]. The emergence of such complexity in the intertwined nature of eco-evolutionary dynamics is a natural outcome of biological processes derived from mechanistic first principles [DIS17]. Still, in the limit of weak selection, the qualitative behaviour of the fixation process of a mutant trait is captured by the corresponding replicator dynamics of evolutionary games yielding a clean separation of ecological and evolutionary scales.

## Acknowledgements

The authors appreciate generous funding from the Max Planck Society.

## A The model and its diffusion approximation

We base our model on a stochastic competitive Lotka-Volterra system which can be derived from the following microscopic reactions:

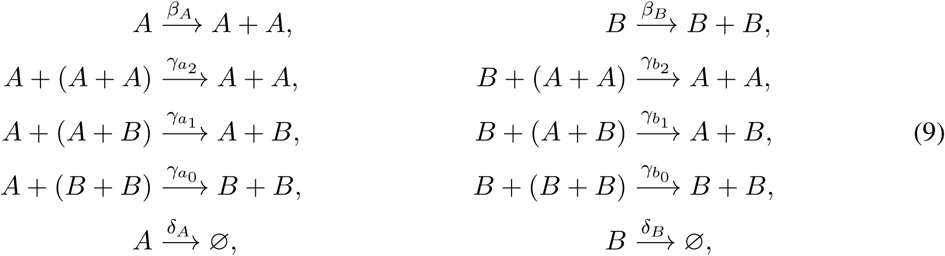

The competition reaction rates take the form 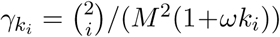, where *M* is a scaling parameter describing the size of the population in stationarity. The parameters *k*_*i*_ are interpreted as entries of a payoff matrix for a three player game, i.e.

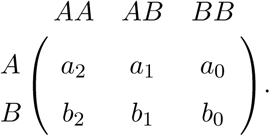

The binomial factor corrects according to the combinatorial possibilities of the corresponding interaction as done in [MN04], i.e. it counts the possible combinations of encountering two individuals of the types participating in the interaction. We note that this combinatorial correction is not the same as the often assumed mass-action kinetics used in studying chemical reactions [Gil76, AK15]. However, it can be translated into that classical setting by adjusting the corresponding reaction rates. Lastly, the factor *M* ^2^ is scaling the interaction rates such that in equilibrium the population indeed is of order *M*. Thus, the reactions are proportional to the type densities rather than their absolute number as is standard in Lotka-Volterra dynamics.

We proceed by writing down the transition rates to go from a state **n** = (*n*_*A*_, *n*_*B*_) to another accessible state. These rates read

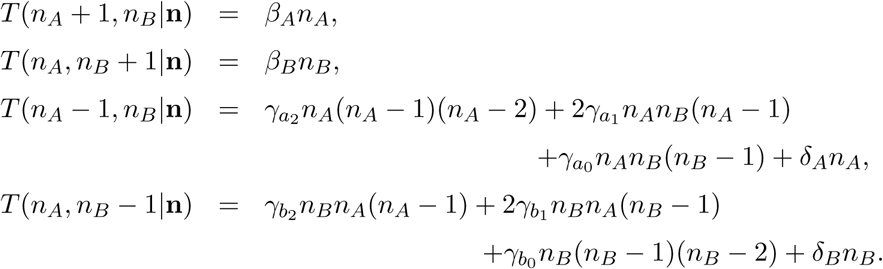

Using these rates we write down the infinitesimal change of the stochastic system. In mathematical terms this is the infinitesimal g enerator *G* o f t he M arkov p rocess c orresponding t o t he microscopic reactions in (9). Thus, for a twice differentiable function *f* we find (just considering the reactions affecting type *A*)

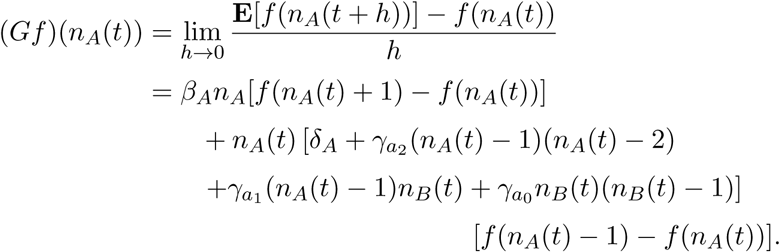

Setting 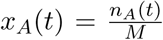, expanding in terms of 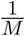 and neglecting terms of order higher than *M*^*–*1^ we obtain

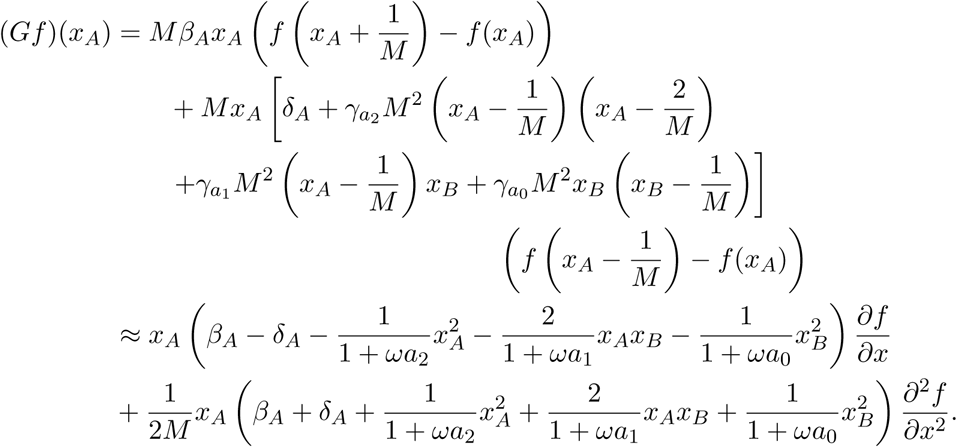

Letting *M* tend to infinity we find the deterministic approximation, see [EK86, Theorem 4.8.2, Corollary 4.8.6] or [Ace02] 19.25]kallenberg:book:2002:

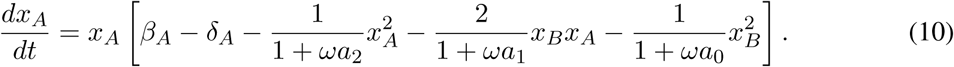

Similarly for trait B we have

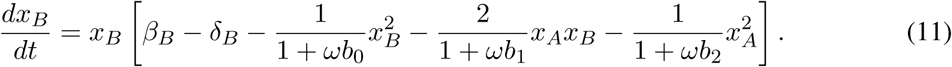

Additionally, we find that the infinitesimal generator of the whole system corresponds to a set of stochastic differential equations given in equation (3) in the main text (cf. [Kal02, Chapter 21]).

For completeness we write down the infinitesimal generator corresponding to the two-dimensional stochastic differential equation:

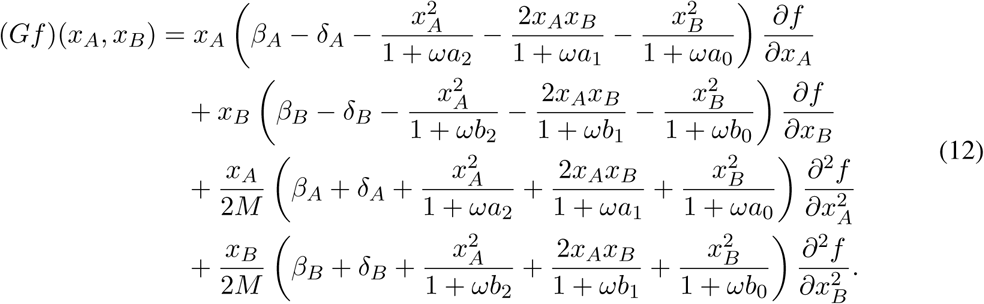

## B Approximating the fixation probability

To get an approximation for the fixation probability of type *A* individuals we first transform the system to the parameter space 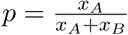, fraction of type *A* particles, and *z* = *x*_*A*_ + *x*_*B*_, the population size. Additionally, setting *β*_*A*_ = *β*_*B*_ = *β* and *δ*_*A*_ = *δ*_*B*_ = *δ* the transformed generator is then given by

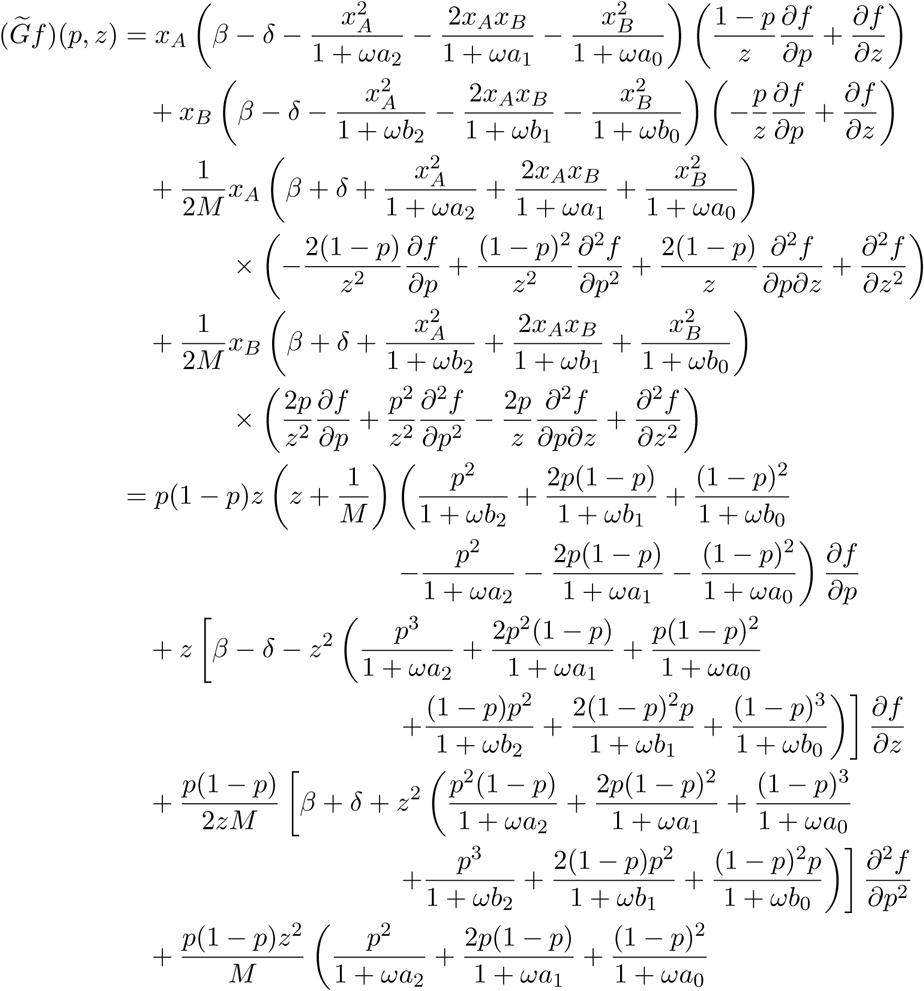

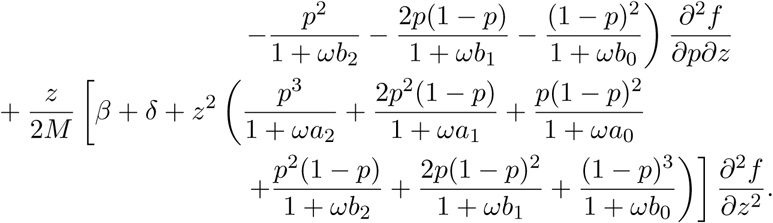

Again, as done in the previous section, we can derive the limit for *M → ∞* and obtain the deterministic evolution of *p* and *z* which reads as

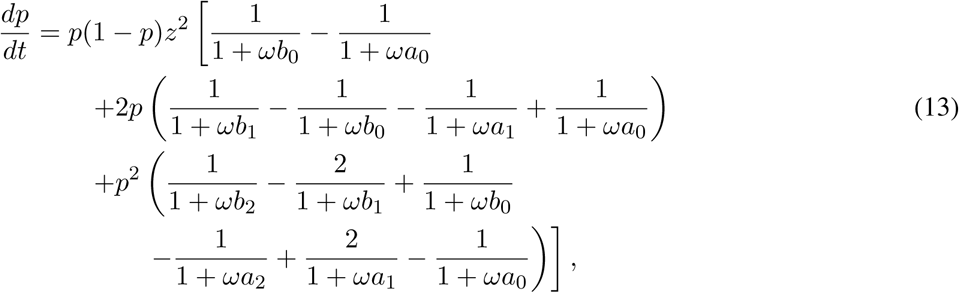

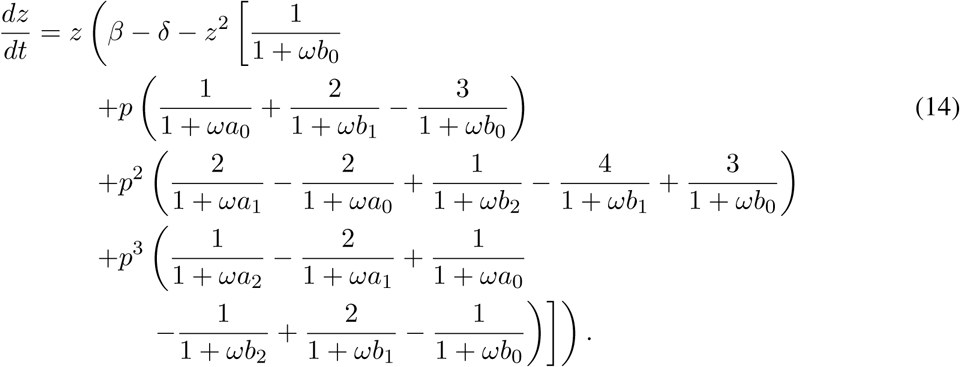

Solving for the fixed point in the frequency space *p*, i.e. 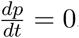, we obtain the following fixed points:

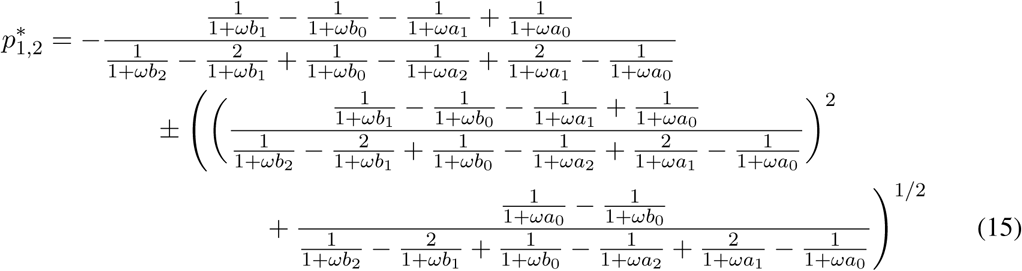

In a next step we assume weak selection, i.e. *ω ≪* 1 which means that the payoffs only have a small impact on the overall evolution of the model. This allows us to simplify the generator and we find

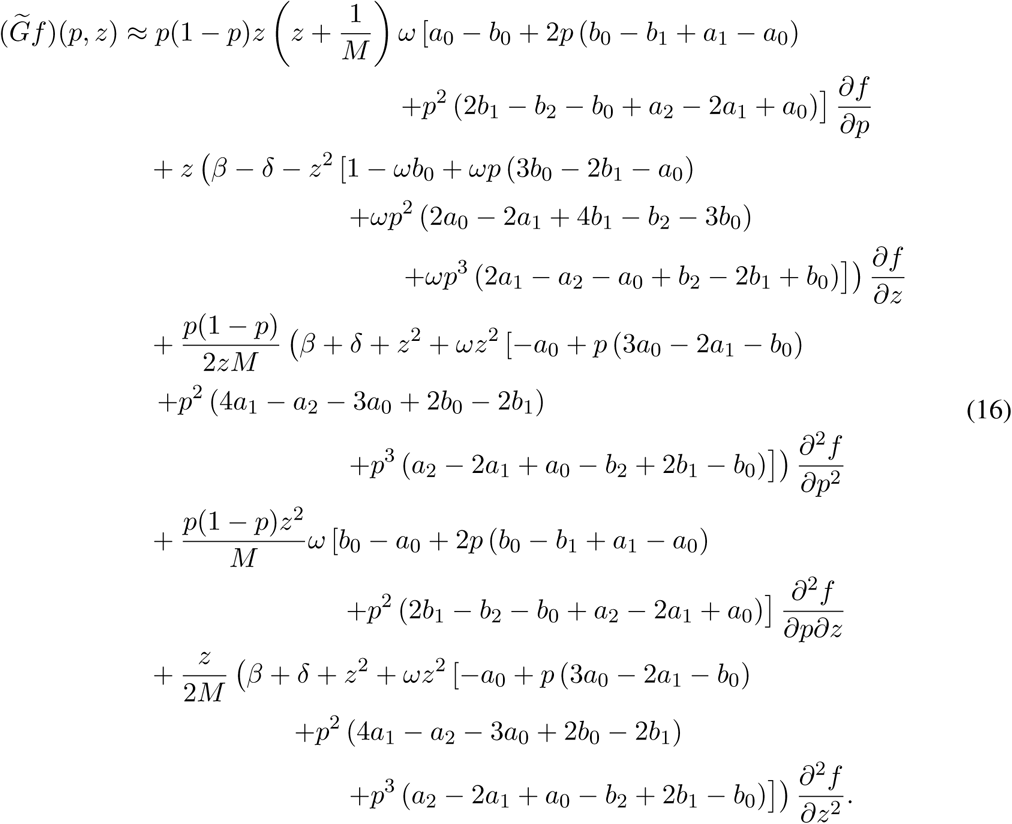

Using the same techniques as developed in [Lam06] and refined for this setting in [CT18, Appendix B, C, D] we find the following approximation for the fixation probability:

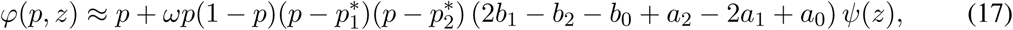

where in the limit of weak selection the internal equilibria 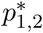 are given by (see also equation (15))

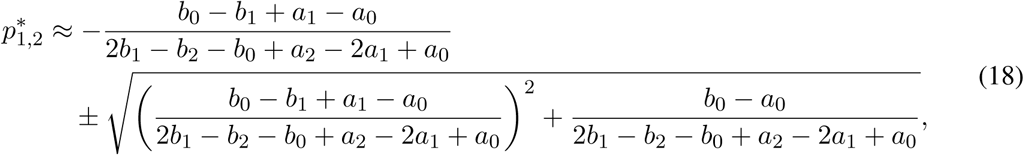

To see that the formula for the fixation probability holds we plug in *ϕ* into 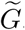. Setting 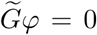 and simplifying we end up with an equation for *ψ*(*z*):

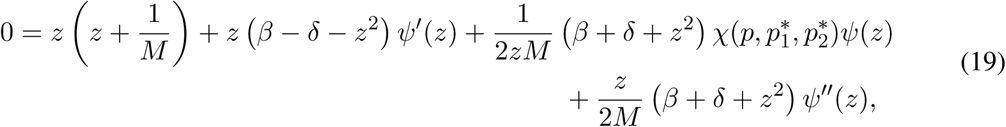

where

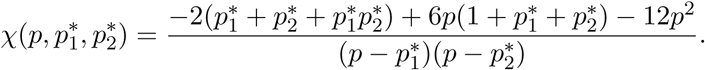

For the method to work it is important that *ψ* depends only on the total population size *z*. Hence, for ψ to be independent of the frequency of mutants *p*, we need *χ* = constant which can be derived from

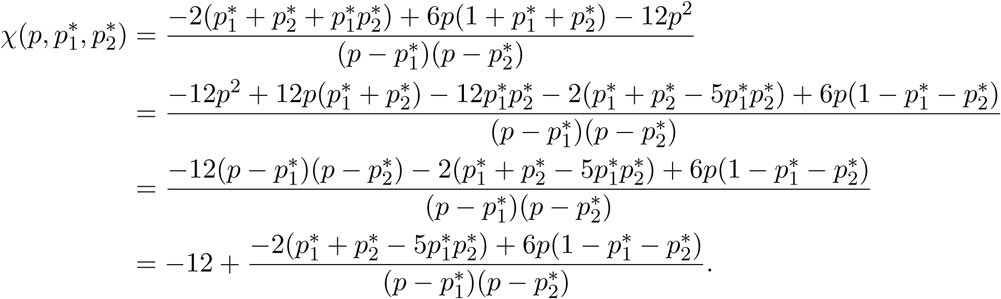

For the second term to vanish (or at least to be negligible) we find the following conditions for the fixed points:

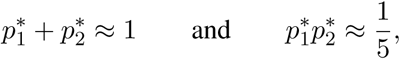

which in the end gives 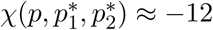. Inserting this into equation (19) yields

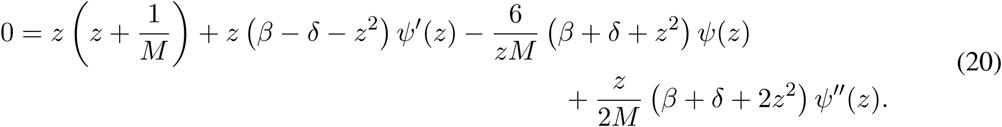

### B.1 One interior fixed point

In this section we assume that the payoff matrix allows for one internal equilibrium close to 1*/*2 and further satisfies

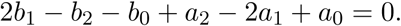

Inserting this into equation (16) we find

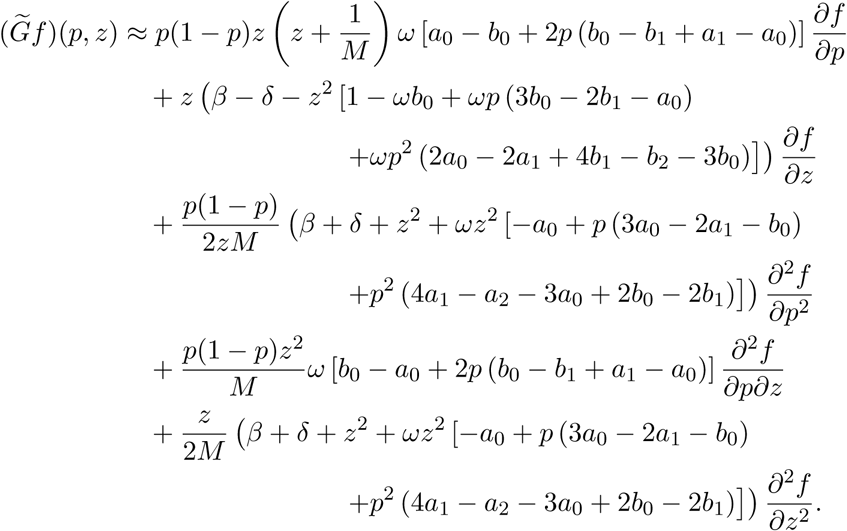

Setting

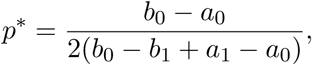

we find that for *ω ≪* 1 the equality 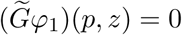 is satisfied for

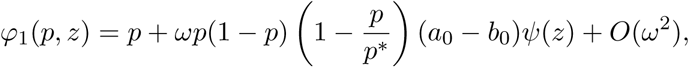

with *ψ*(*z*) solving

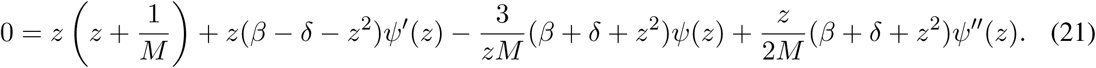

### B.2 No interior fixed point

In this section we assume that either *a*_*i*_ *> b*_*i*_ or *b*_*i*_ *> a*_*i*_ for all *i* = 0, 1, 2. This implies that there is no internal fixed point in the associated Lotka-Volterra system, i.e. one of the two strains is strictly dominating. We first rewrite equation (16) as follows

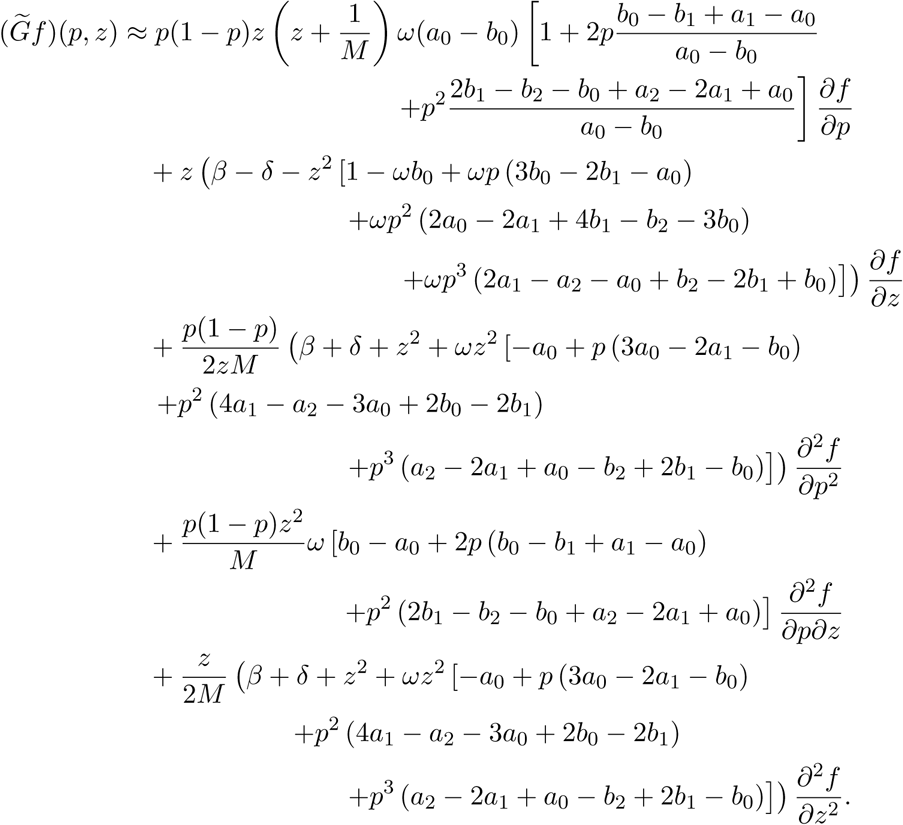

Further assuming that the fractions 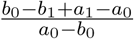 and 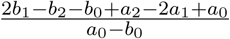 are close to 0, i.e. the product with *ω* is negligible, we find that

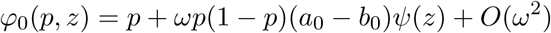

solves 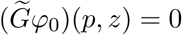 with *ψ*(*z*) solving

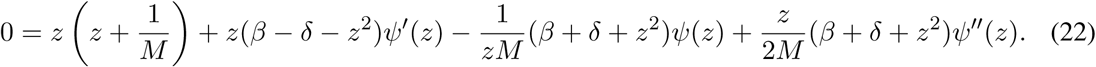

### B.3 Extension to *d* players

We now extend the formalism to the general setting with *d–*players and two strategies. The payoff matrix is then given by

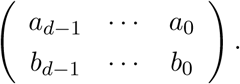

The death rates due to competition need to change accordingly, i.e. a death of a type *A* particle occurring due to the interaction with *k –* 1 type *A* and *d – k* type *B* individuals is given by

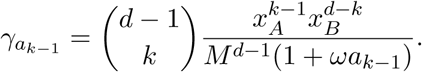

Instead of scaling these interactions with 1*/M* ^2^ we need the scaling 1*/M*^*d–*1^ in order to obtain a reasonable diffusion limit. Performing the same steps as in the three player case we end up with the following stochastic differential equations:

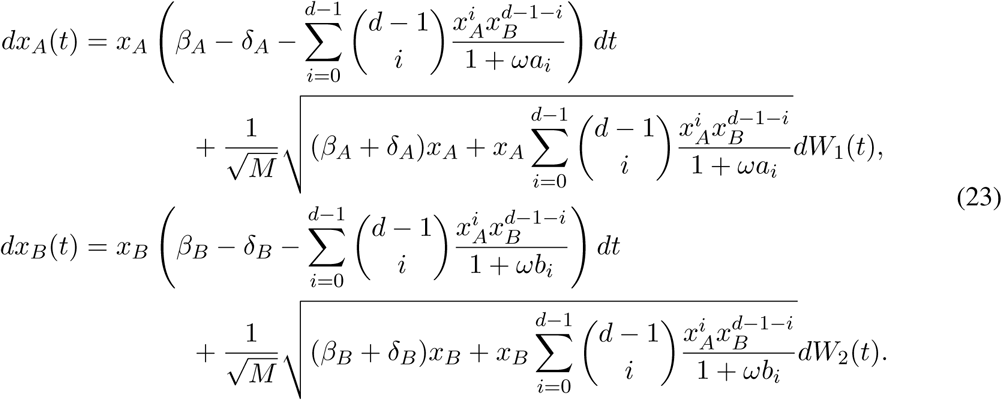

The transformed generator, i.e. in terms of *p* and *z*, can again be derived analogously to the three player scenario and reads:

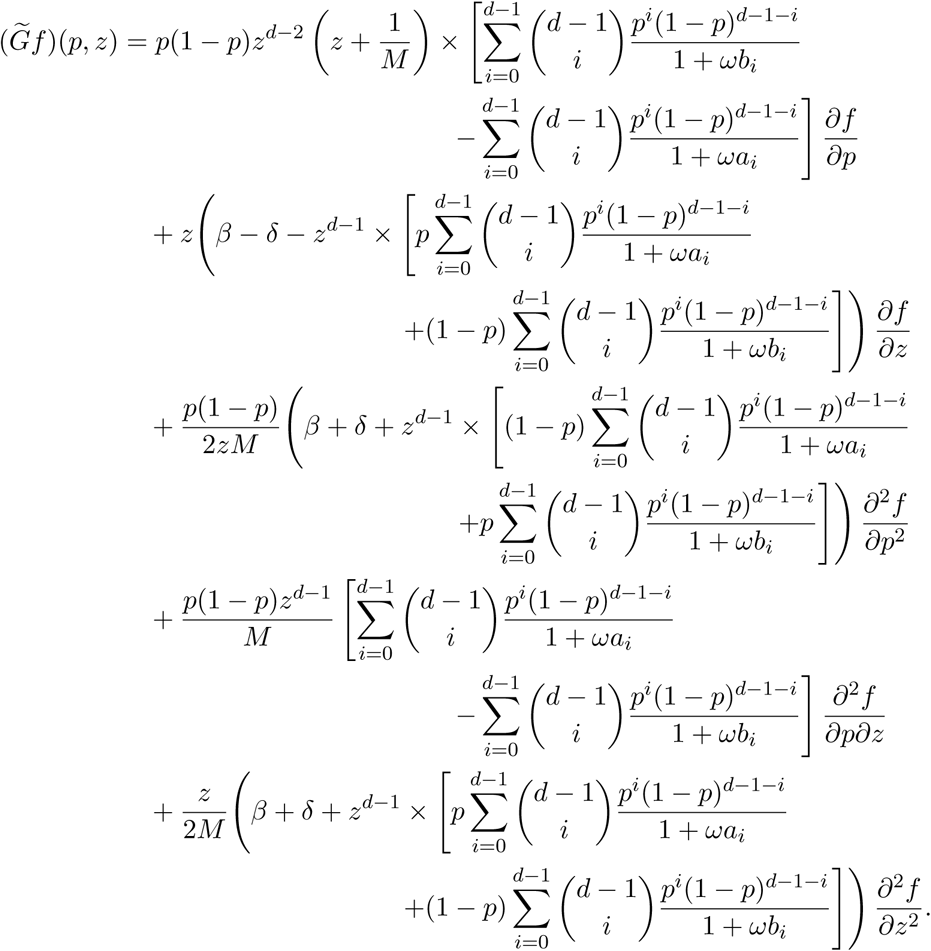

The solution to 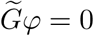 for *ω ≪* 1 can be approximated by

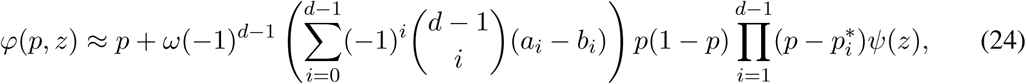

where 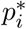 are the roots of

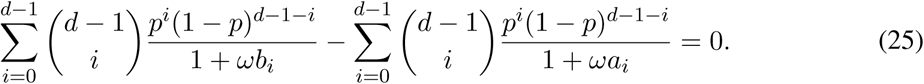

This can be seen by plugging in the resulting approximation of *ϕ* into 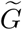. Along the way one needs to see that 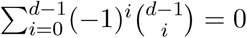 which can be proved by induction. Then the remaining equation that *ψ* needs to satisfy is given by

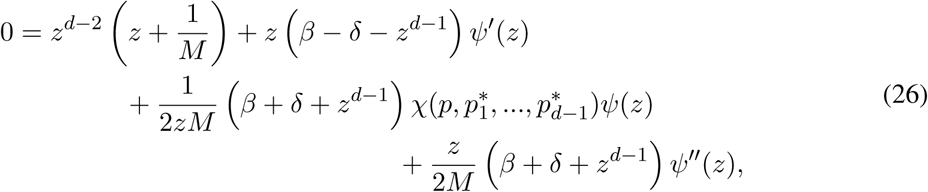

where

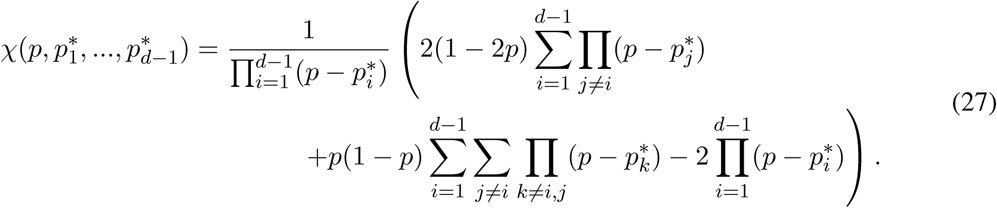

For *χ* to be independent of *p* this formula gives conditions on the location of the fixed points 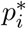 such that the above approximation of the fixation probability, i.e. the separation of the *p* and *z* coordinates, is valid. However, the general form of this condition is beyond the scope of our analysis.

Following the same reasoning as above (before equation (20)) and just gathering the terms of order *p*^*d*-1^ we get

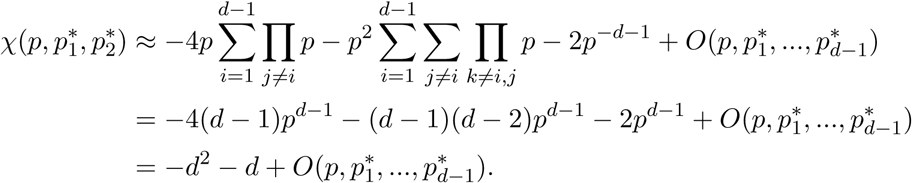

Assuming *χ* to be independent of *p* would then give the following differential equation for *ψ*(*z*):

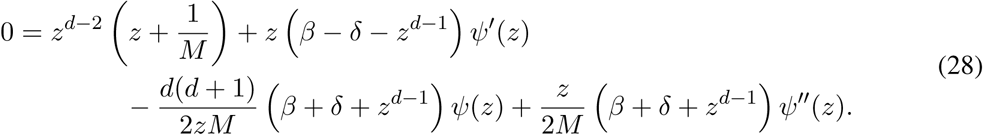

### B.4 Numerical evaluation of *ψ*

To calculate values of *ψ* we numerically evaluate, dependent on the number of players and the number of internal equilibria, equation (20), (21), (22) or (28), respectively. We use the predefined function “`solve bvp`” from the `scipy.integrate` library in Python, [JOP+]. We need to provide boundary values for the algorithm to work with. In particular we evaluate *ψ* in the interval [10^*–*4^, 10] with boundary values *ψ*(10^*–*4^) = 10^*–*4^ and *ψ*(10) = ln(10). The concrete choice of the boundary values is not relevant since the method is quite robust. For instance, we tested for different values of *ψ*(10^*–*4^) *∈* [10^*–*8^, 0.1] and all solutions gave the same values for *z* = 2. The same applies to the boundary value *ψ*(10). For a more thorough analysis of *ψ* we refer to [CT18, Appendices E,F].

### B.5 Fixation probability under strong selection

We provide an approximation of the fixation probability in the *d*-player eco-evolutionary model in case of the snowdrift game dynamics (for details see the main text, Section 3.4). As in Figure 6 in the main text, we vary the selection intensity *ω* and analyse the fixation behaviour of the cooperative trait. The dynamics allows for two internal fixed points 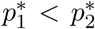 where 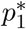 is locally unstable and 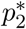 is locally stable.

Deviating from the main text, we start the simulations in the close vicinity of the fixed point 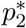. Weak selection then predicts fixation probabilities close to the initial frequency of cooperative individuals, i.e. 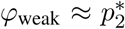. As we see in Figure 7 this is a good approximation for the simulation outcome until some intermediate value of *ω*.

**Figure 7:**
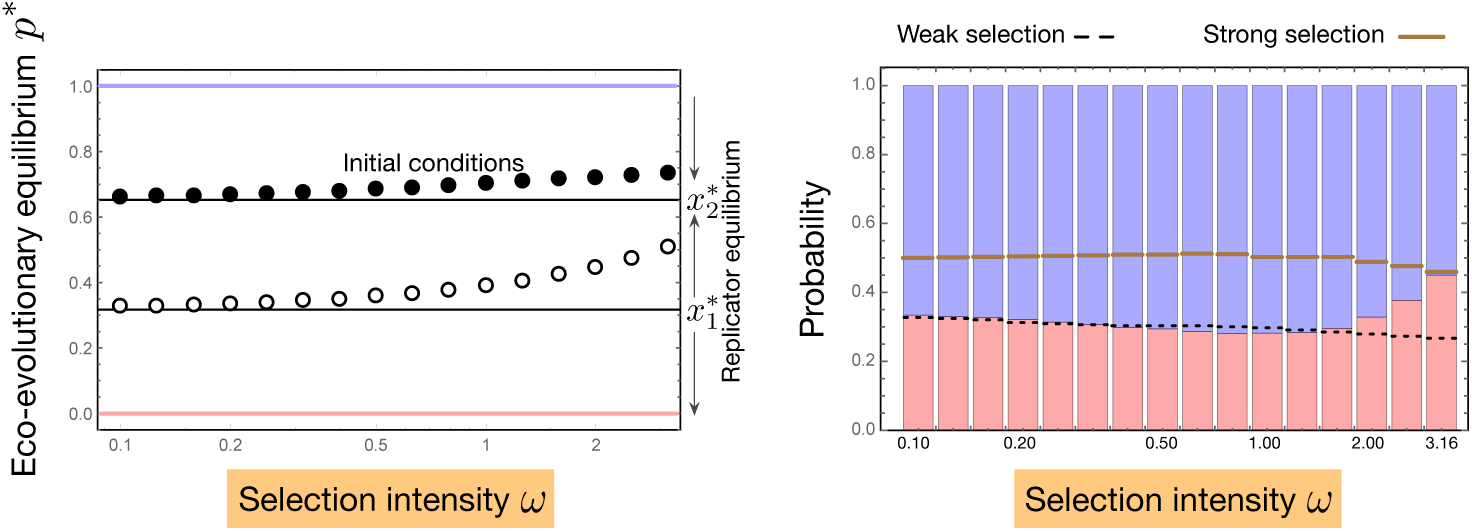
Comparison between weak and strong selection. Re-evaluating the findings from Figure 6 (where the initial condition was set to *p*_0_ = 0.5) we now consider 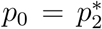 (filled circle). The deterministic behaviour of the system is visualized in the left panel. By varying the selection intensity *ω* we also alter the location of the deterministic fixed points. For weak selection the effect of the selection gradient is minimal and drift plays a dominant role and hence the fixation probabilities are close to neutral (dashed lines in the right panel). As selection increases, the stable fixed point, close to all *C*, attracts and holds most of the trajectories. This is why we describe the escape dynamics of the attractor region by a Brownian motion, leading to equation (29). These values are shown as solid lines in the right panel. They are indeed approaching the simulation results for stronger selection intensities. In these cases the trajectories either hit the monomorphic *C* state or overcome the unstable equilibrium 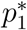 (empty circle) and end in the monomorphic *D* state. We have performed 10^5^ Gillespie simulations for *M* = 100 and a 20 player snowdrift game with an initial population of 200, *b* = 1.5, *c* = 1, *β* = 0.6, *δ* = 0.1, *ψ ≈* 0.53 and 1 + *ω*(*matrix*) mapping for different intensities of selection. The bar charts show the probability of a trajectory fixing in either *allC* (blue) or *allD* (red). All simulations were executed until one of the types fixed. The selection intensity ranges as *ω* = 10^*–*1.0^,…, 10^0.5^.

For very strong selection we argue that the system is dominated by the replicator dynamics. Hence, trajectories stay close to 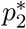 for a long time and randomly fluctuate around this stable equilibrium. The fixation probability is then given by the escape probability of the attracting domain at the boundary *p* = 1 which can be calculated by stochastic diffusion theory (see for instance [Ewe04, Chapter 4], i.e.

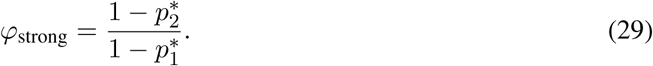

For the strongest considered selection intensity this exactly fits the simulation result (right-most value in the right subfigure of Figure 7) supporting our heuristic reasoning in the main text.

This has an intriguing implication – both weak and strong selection can be explained by random drift. Still there is a difference since in the weak selection limit the replicator dynamics is close to neutral, i.e. the stability of the fixed points just has a minor impact on the dynamical behaviour. Hence, the fixation process can be described by a Brownian motion over the whole frequency space resulting in a fixation probability close to the initial frequency. For strong selection however, the deterministic dynamics defines the trajectories of the individual based model. Thus, in this case the escape behaviour out of the attractor region of the stable fixed p oint i s d escribed b y a B rownian m otion y ielding the approximation obtained in equation (29).

